# Mapping of the Classical Mutation *rosette* Highlights a Role for Calcium in Wound-induced Rooting

**DOI:** 10.1101/2022.09.24.509321

**Authors:** Abelardo Modrego, Taras Pasternak, Moutasem Omary, Alfonso Albacete, Antonio Cano, José Manuel Pérez-Pérez, Idan Efroni

**Affiliations:** The Institute of Plant Sciences and Genetics in Agriculture, Faculty of Agriculture, The Hebrew University, Israel; Instituto de Bioingeniería, Universidad Miguel Hernández, 03202 Elche, Spain; Departamento de Nutrición Vegetal, CEBAS-CSIC, 30100 Murcia, Spain; Departamento de Biología Vegetal (Fisiología Vegetal), Universidad de Murcia, Murcia, Spain

**Keywords:** Arabidopsis, auxin transport, BIG, calcium, tomato, UBR4, wound-induced roots

## Abstract

Removal of the root system induces the formation of new roots from the remaining shoot. This process is primarily controlled by the phytohormone auxin which interacts with other signals in a yet unresolved manner. Here, we study the classical tomato mutation *rosette* (*ro*) which lacks shoot-borne roots. *ro* plants were severely inhibited in the formation of wound-induced roots and have reduced rates of auxin transport. We mapped *ro* to the tomato ortholog of the *Arabidopsis thaliana BIG* and the mammalians *UBR4/p600.* RO/BIG is a large protein of unknown biochemical function. In *A. thaliana, BIG* was implicated in the regulation of auxin transport and calcium homeostasis. We show that exogenous calcium inhibits wound-induced root formation in both tomato and *A. thaliana ro/big* mutants. Exogenous calcium antagonized the root-promoting effects of the auxin IAA, but not of 2,4-D, an auxin analog that is not recognized by the polar transport machinery, and accumulation of the auxin transporter PIN1 was sensitive to calcium levels in the *ro/big* mutants. Consistent with a role for calcium in mediating auxin transport, both *ro/big* mutants and calcium-treated wild-type plants were hypersensitive to treatment with polar auxin transport inhibitors. Subcellular localization of BIG suggests that like its mammalian ortholog, it is associated with the endoplasmic reticulum (ER). Analysis of subcellular morphology revealed that *ro/big* mutants exhibited disruption in cytoplasmic streaming. We suggest that RO/BIG maintain auxin flow by stabilizing PIN membrane localization, possibly by attenuating the inhibitory effect of Ca^2+^ on cytoplasmic streaming.

## Introduction

Plants can recover from loss or damage to their root system by the production of new roots from their shoots and hypocotyls. These roots are generally referred to as adventitious roots (AR), but in specific developmental contexts called shoot-borne-roots (SBR), and when formed in response to injury, are called wound-induced roots (WiR). WiR initiation is regulated by a range of phytohormones which play a role in all stages of root formation (Bellini *et al*., 2014; Steffens & Rasmussen, 2016; Omary *et al*., 2022).

Auxin (Indole-3-Acetic-Acid; IAA) is the central hormone controlling WiR initiation (Pacurar *et al*., 2014), and increased levels of endogenous or exogenous auxin promote root formation in many plant species, a property commonly utilized for clonal propagation (Boerjan *et al*., 1995; Delarue *et al*., 1998; de Klerk *et al*., 1999; Druege *et al*., 2016; Lakehal & Bellini, 2018; Guan *et al*., 2019). The transcriptional response to auxin is mediated by AUXIN RESPONSE FACTOR transcription factors which, in the presence of auxin, are released from their inhibitors, AUX/IAAs, and allowed to bind auxin response elements in promoters of target genes (Weijers & Wagner, 2016). Consistent with auxin’s role in WiR initiation, the synthetic auxin-responsive promoter DR5 (Ulmasov *et al*., 1997) is induced at the earliest stages of WiR initiation (Welander *et al*., 2014; Della Rovere *et al*., 2016).

Auxin distribution within tissues is determined both by its site of biosynthesis and by the activity of polar and non-polar auxin transporters. Auxin is produced from the amino acid tryptophan by the enzymes TRYPTOPHAN AMINOTRANSFERASE OF ARABIDOPSIS and YUCCA (Zhao, 2014). The cellular uptake of auxin is mediated by the AUXIN-RESISTANT1 (AUX1)/LIKE AUX1 (AUX1/LAX) family of non-polar transporters (Swarup & Péret, 2012) while its export is mainly controlled by the PIN-FORMED transporters (PINs), which are polarly localized on the plasma membrane (Adamowski & Friml, 2015). While most PINs are localized to the plasma membrane, the *Arabidopsis (Arabidopsis thaliana)* PIN5, PIN6, and PIN8 were shown to also localize to the endoplasmic reticulum (ER) membrane, where they regulate intracellular auxin distributions together with the PIN-LIKE proteins (Adamowski & Friml, 2015; Simon *et al*., 2016; Sauer & Kleine-Vehn, 2019).

The PINs are required for directional auxin transport within plant tissues and disruption of their activity, either genetically or using the chemical inhibitor naphthylphthalamic acid (NPA), results in altered auxin distribution within tissues (Sabatini *et al*., 1999; Adamowski & Friml, 2015; Abas *et al*., 2021). NPA blocks IAA transport by directly binding the PIN proteins (Abas *et al*., 2021; Teale *et al*., 2021; Ung *et al*., 2022), but other possible NPA targets were suggested, such as ATP-binding cassette transporters or their interacting partner TWISTED DWARF (Teale & Palme, 2018). An early putative NPA target is the large callosin-like protein BIG/TRANSPORT INHIBITOR RESPONSE 3 (TIR3) which plays a role in the regulation of the auxin transport. *big* mutants have reduced rates of auxin transport, show reduced binding of NPA, and reduced levels of PIN proteins (Ruegger *et al*., 1997; Gil *et al*., 2001; Liu *et al*.,2022).

How endogenous auxin gets to the initiating WiR is not entirely known. Mutants defective in auxin import or PIN-mediated auxin export have reduced number of WiR (Sukumar *et al*., 2013; Lee *et al*., 2019), and using local application of NPA to block auxin transport from the shoot results in a reduction in the number of formed roots. This suggests that most of the auxin required for WiR formation is shoot-derived (Ahkami *et al*., 2013; Sukumar *et al*., 2013; Alaguero-Cordovilla *et al*., 2021).

Apart from hormones, other molecules also mediate WiR initiation. Amongst these, Ca^2+^ is a universal secondary messenger involved in many cellular processes, including auxin signaling (Dodd *et al*., 2010; Vanneste & Friml, 2013). Ca^2+^ can be sensed by calmodulin (CaM), a small protein that regulates the activity of other proteins by interacting with their calmodulin-binding domain (Dodd *et al*., 2010). The effects of Ca^2+^ levels on WiR initiation are complex and were mostly studied in tissue culture conditions.

Ca^2+^ treatment can enhance the rooting promoting effect of IAA in poplar (Bellamine *et al*., 1998) and mung bean *(Phaseolus aureus)* when applied together with borate (Jarvis & Yasmin, 1985), but had no effect in *Dalbergia* (Ansari & Kumar, 1994). Chelating Ca^2+^ using EGTA partially inhibited adventitious rooting in poplar and mung bean (Bellamine *et al*., 1998; Li & Xue, 2010) but a transient EGTA treatment promoted rooting in sunflower (Kalra & Bhatia, 1998). In cucumber, inhibition of calmodulin activity counteracted the root promoting effects of exogenous auxin or nitric oxide (Lanteri *et al*., 2006; Niu *et al*.,2017) and application of Ca^2+^ activated ethylene-signaling to relieve the repression of rooting promoted by salt stress (Yu *et al*., 2019). In mung bean, it was suggested that Ca^2+^ acts downstream of auxin, H_2_O_2_, and nitric oxide (Lanteri *et al*., 2006; Li & Xue, 2010). Overall, how Ca^2+^ and auxin interact during WiR initiation is still unclear.

Tomato *(Solanum lycopersicum)* has been used as a model species for developmental studies of both shoot and root systems and recently also as a model for WiR formation (Alaguero-Cordovilla *et al*., 2021). Several mutations that affect shoot-borne roots production in tomato were identified, and those serve as good candidates for WiR regulators (Zobel, 1975). The classical tomato mutation *rosette (ro)* displays severe reduction in internode elongation, reduced branching, and complete sterility. Notably, these plants were described to have no SBR and a reduced number of lateral roots (LRs; Butler, 1954; Zobel, 1975; Schiefeibein *et al*., 1991). Although first described in 1954, the causal mutation underlying the *ro* phenotype was never identified. Here, we show that *ro* mutants are defective in WiR initiation. We map *ro* to the tomato ortholog of *BIG* and show that the reduction in WiR formation in these mutants is likely attributed to Ca^2+^-related disruption of the polar auxin transport machinery. Finally, we show that RO/BIG is likely localized to the ER and is important for maintaining proper cytoplasmic streaming, suggesting a possible link between subcellular organization and tissue-wide auxin distribution patterns.

## Results

### *rosette* is defective in wound-induced root production and auxin transport

Tomato *ro* mutants are dwarf, exhibit little internode elongation and their stems and hypocotyls are narrow compared to wild-type (WT) plants (Fig. 1A-C; Butler, 1954). Dissection of the hypocotyl of four-week-old plants revealed that both the WT and *ro* mutants had a similar organization with 4 opposing vascular bundles. The width of the cortex was reduced by 30% in the *ro* mutant (WT 578±7.6 μm, *ro* 403±10.2 μm; n=8; p<0.0001; Student’s t-test), but most of the reduction in stem width was due to a 75% reduction in stele width (WT 2123±198 μm, *ro* 528±28 μm; n=8; p<0.0001; Student’s t-test). Additionally, *ro* mutants lacked interfascicular cambium (Fig. 1D-F). As previously reported, uninjured *ro* plants produced no SBR from their hypocotyls (Schiefeibein *et al*., 1991); Fig. 1G, H). To test whether the *ro* mutation also affects WiR production, we removed the root system of four-week-old *ro* plants and WT siblings and allow the cuttings to root in deionized water. After 10 days, wild-type plants formed a large number of roots all along the hypocotyl while *ro* cuttings formed only a few roots at the base of the hypocotyl (WT 54±13, *ro* 5.1±2.8 roots per plant; n=20; p<0.0001; Student’s t-test; Fig. 1I, J).

**Fig. 1.**
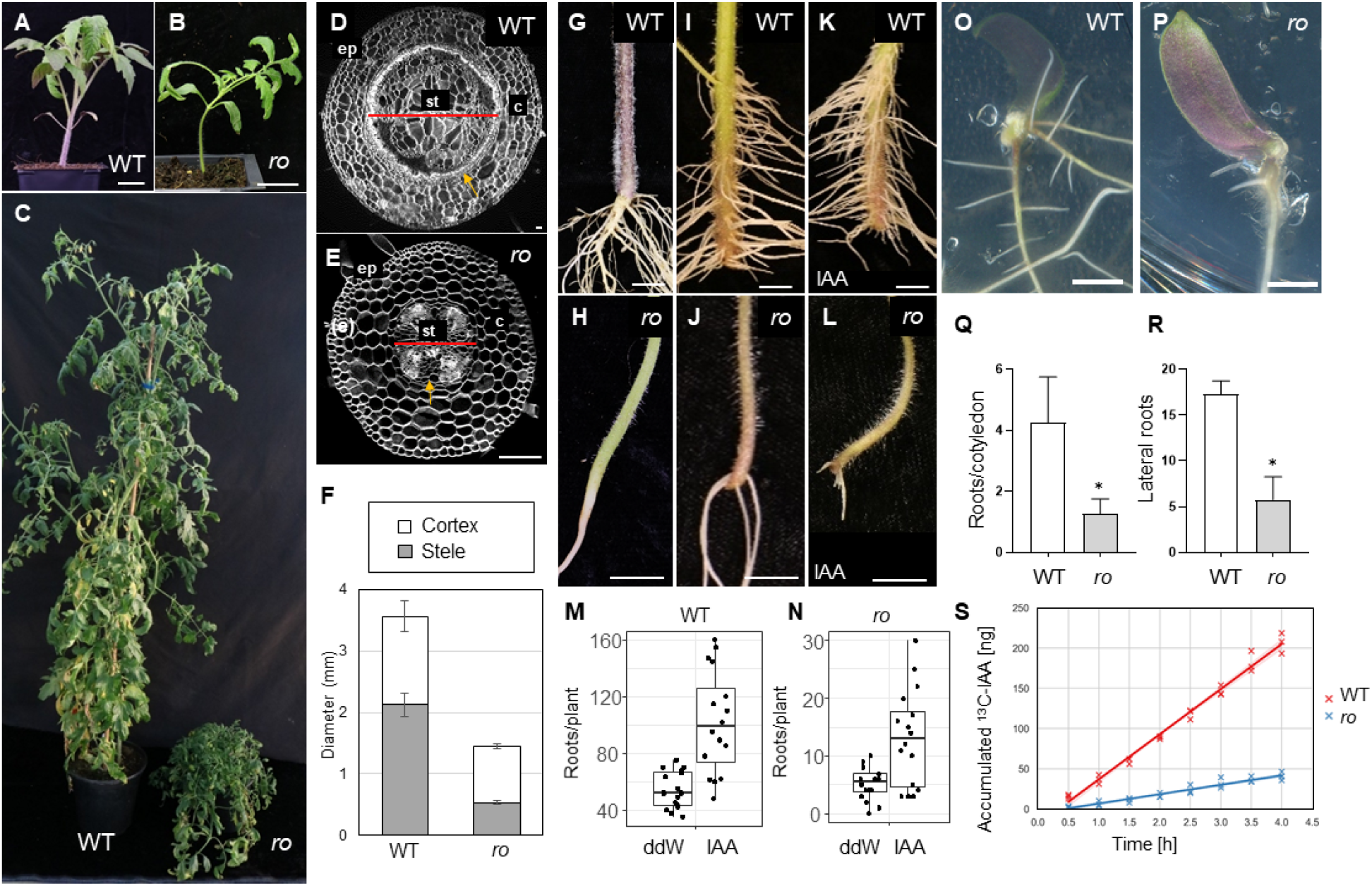
The classical *rosette* mutation is defective in wound-induced root production. (A-C) Tomato WT and *rosette* (*ro*) one-month months (A-B) and five-month-old (C) plants. (D and E) Confocal images of hypocotyl cross sections of WT (D) and *ro* (E). Cell wall was stained with SCRI Renaissance 2200. Note that the WT and *ro* panels are at different scales. ep-epidermis; st – stele; c – cortex. Red line marks the diameter of the stele. Orange arrows points to the interfascicular cambium in (D) and its absence in (E). (F) Diameters of entire hypocotyl and its stele in WT and *ro* mutants (n=8 for WT and *ro*). (G and H) Close up images of hypocotyls of WT (G) and *ro* (H). (I-L) WiR in hypocotyls production 10 days after cutting (dac) in WT (G and I) and *ro* (H and J), either when in incubated in water (I and J) or with 10 μM IAA (K and L). (M-N) Quantification of WiR production in WT (M) and *ro* (N) cuttings incubated in water or 10 μM IAA. Bar indicates the mean. Two-tailed Student’s t-tests: WT-ddW Vs IAA, p<0.01 (n=20 and 20 for ddW and IAA respectively); *ro*-ddW Vs IAA, p<0.01 (n= 20 and 20 for ddW and IAA respectively). (O-P) Cut cotyledons of 7-8 days old WT (M) and *ro* (N) plants on TK4 media with 50 nM NAA for 12 days. (Q-R) Quantification of WiR (Q) and LR (R) number formed on cotyledons (n=4 for each condition; p<0.01, Two tailed Student’s t-test). (S) Accumulated transported IAA in WT and *ro* tomato hypocotyls. n=3 for each genotype. Scale bars are 1 cm in (A-B, G-L, O-P) and 100 μm in (D and E).

To test whether exogenous auxin can rescue the *ro* WiR phenotype, we treated cuttings with 10 μM IAA. As previously reported (Alaguero-Cordovilla *et al*., 2021), this treatment increased the number of roots in WT plants. While IAA also increased the number of WiR in *ro* plants, the response was attenuated, the number of roots remained low and they were confined to the bottom of the hypocotyl, near the cut site (Fig. 1K-L, M-N). We further evaluated whether auxin-induced *de novo* root formation was affected by the *ro* mutation using young cotyledon explants. When tomato cotyledons are dissected and placed on agar media containing the synthetic auxin NAA, they can produce AR. *ro* mutants developed fewer roots and these produced fewer LR than WT (Fig. 1O-R).

Previously, it was shown that inhibition of auxin transport rates using NPA reduced the number of roots formed on cut tomato hypocotyl (Alaguero-Cordovilla *et al*., 2021). To test whether auxin transport rates are affected in *ro*, we applied radiolabeled IAA to the apical side of hypocotyl sections of WT and *ro* mutants and detected its accumulation in the basal side. We found that IAA transport intensity through the hypocotyl was significantly reduced in *ro* mutants, with a five-fold difference in IAA transport rates compared to WT (Fig. 1S; 56.03±3.61 ng h^-1^ in WT and 11.64±0.97 ng h^-1^ in *ro* mutants; p-value <0.001; Student’s t-test). Overall, our results suggest that *ro* is disrupted in the formation of WiR, possibly due to the inability to form sufficient auxin concentrations at the hypocotyl\cotyledon base.

### *rosette* is a mutation in the tomato ortholog of *BIG*

To better understand the role of *RO* in WiR initiation we turned to mapping-by-sequencing to identify the causal mutation. Previous genetic linkage experiments have mapped *ro* to the long arm of chromosome 2, in a linkage group with *dwarf* (*Solyc02g089160;* Butler, 1954). We re-sequenced a pool of 12 *ro* mutant plants and 12 WT-looking siblings to a coverage of x40 and identified a region of homozygosity in the *ro* mutant pool on chromosome 2 between 50,475,714bp and 51,141,636 bp (ITAG2.5 coordinates). This region contained 88 genes, including the *DWARF* locus. Only one mutation that disrupted the amino acid sequence was identified in this region; a transversion causing a premature stop at the 8^th^ exon of *Solyc02g089260* (11725A>T; Fig. 2A; Supplementary Fig. S1A-B). To confirm that the *ro* mutant is caused by a disruption to *Solyc02g089260*, we generated CRISPR-edited plants of this gene in tomato M82 background. A CRISPR-induced insertion at the 3rd exon caused an early frameshift, likely resulting in a null allele (Fig. 2A; Supplementary Fig. S1A). These mutant plants were similar to the *ro* plants, had no internode elongation, their leaves were small and simple, and they lacked SBR. As in *ro* plants, *Solyc02g089260^RISPR^* mutants could form a few WiR when the root system was removed, but those were confined to the base of the hypocotyl (Fig. 2B-D). To confirm that the *ro* phenotype is caused by disruption of *Solyc02g089260*, we performed an allelism test and crossed *ro/+* with *Solyc02g089260^CRISPR^/+* plants. About ¼ of the F1 progeny had the *ro* phenotype, indicating that *ro* is allelic to *Solyc02g089260^CRISPR^* and is likely also a null allele (Supplementary Fig. S1D).

**Fig. 2.**
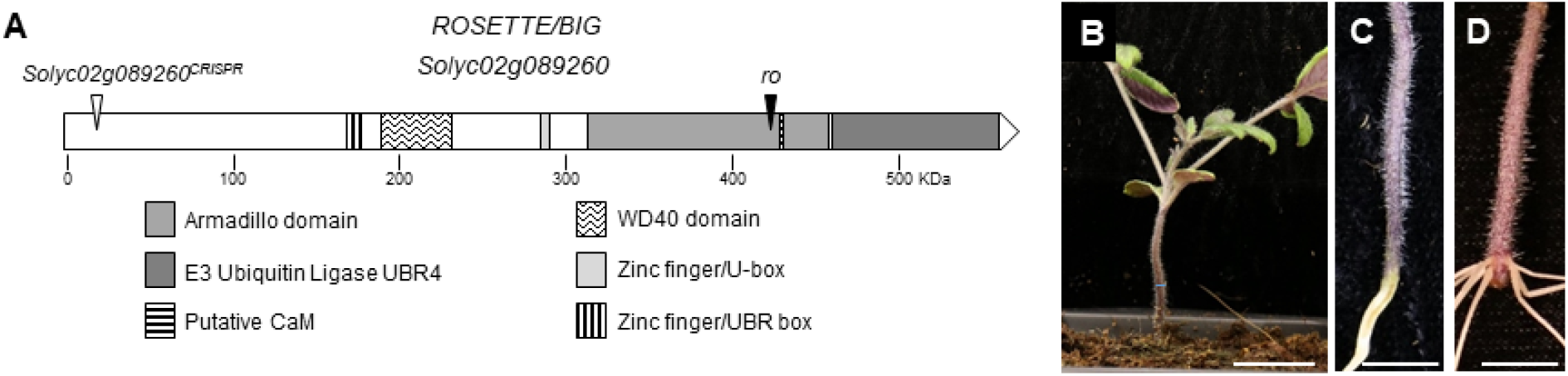
*rosette* maps to the tomato ortholog of *BIG*. (A) Schematic representation of the *RO/BIG* protein. White and black triangles indicate the position of the tomato *Solyc02g089260^CRISPR^* and the *ro* mutations, respectively. (B-D) One-month-old *Solyc02g089260^CRISPR^* mutant plant (B) with no SBR (C) but with WiR (D). Scale bars are 5cm in (B) and 2cm (C-D).

Solyc02g089260 is a very large protein (5104 aa) that has no duplicated family members. Its closest ortholog in *Arabidopsis* is BIG (AT3G02260) and in mammalians, UBR4/p600 (Gil *et al*., 2001; Nakatani *et al*., 2005). Domain annotation using Expasy identified several Zinc fingers, WD40, armadillo, and E3-Ubiquitin Ligase domains. Previous work has shown that BIG also has a non-canonical calmodulin-binding domain (Belzil *et al*., 2013). These domains in *Solyc02g089260* were similar in identity and position to the *Arabidopsis* BIG, indicating that RO, BIG, and UBR4 are likely orthologs (Fig. 2A).

In *Arabidopsis, BIG* was reported to be important for a variety of developmental and physiological processes, such as regulation of light and shade responses, circadian rhythms, gibberellin status, and both root and shoot development (Gil *et al*., 2001; Kanyuka *et al*., 2003; Desgagné-Penix *et al*., 2005; Hearn *et al*., 2018; Liu *et al*., 2022). To determine whether *BIG* plays a similar role in WiR initiation in *Arabidopsis* as it does in tomato, we dissected the root system of five-day-old WT and plants carrying the *tir3-1* allele (which we refer to hereafter as *big*; Supplementary Fig S1A, E; Ruegger *et al*., 1997). Consistent with the *ro* phenotype, the number of WiR in *big* seedlings was significantly reduced and they took longer to appear (Fig. 3A-C). Treatment with IAA increased the number of WiR forming on both WT and *big* hypocotyls (Gutierrez *et al*.,2009) but the number of roots was lower than in comparably treated WT plants (Fig. 3C).

**Fig. 3.**
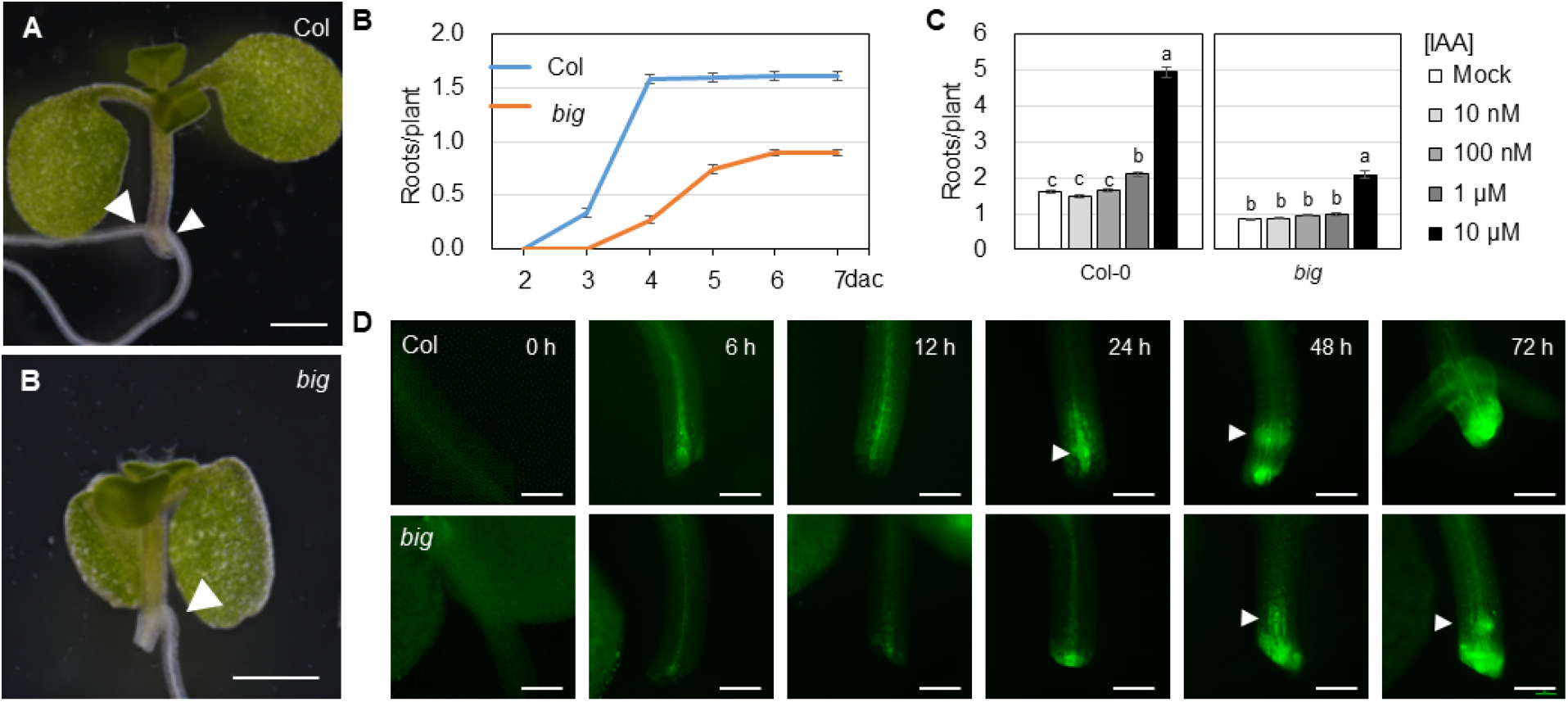
Arabidopsis *big* mutants are defective in wound induced root initiation. (A) *Arabidopsis* Col (WT) and *big* mutant cuttings after 7 days. Arrowheads indicate root primodria. (B) WiR production in Col and *big* cuttings incubated in water (p<0.01, n=124 and 128 for Col and *big* respectively). (C) Number of WiR in *Arabidopsis* Col and *big* mutant cuttings after 7 days with treated with different IAA concentrations (n= 125, 132, 127, 135 and 138 for Col and 137, 130, 131, 137 and 143 for *big* in mock, 10, 100 nM, 1 and 10 μM IAA respectively). (D) Stereo microscope images of *DR5rev:3xVENUS-N7* cuttings in WT (upper panels) and *big* (lower panels) backgrounds. Arrowheads mark root primordia. At least 6 plants were examined and representative images shown. Bars represent the mean ± SE; letters are statistically significant differences (p<0.01; ANOVA and Tukey’s post-hoc test). Scale bars are 1cm (E) and 200 μm (H).

To determine whether root initiation or emergence were defective in *big*, we examined the dynamics of the auxin reporter *DR5rev:3xVENUS-N7*, which marks the early stages of WiR initiation in the hypocotyl (Welander *et al*., 2014). In WT *Arabidopsis*, auxin signaling was induced at the bottom of the hypocotyl at 6h after the cut, and focused signals marking root primordia were apparent at 24h. Auxin signaling was attenuated in *big* mutants and was apparent near the cut site by 24h, while primordia were observed only at 48h-72h (Fig. 3D). This 24h delay in induction of the auxin response in *big*, as compared to WT, is consistent with a similar 24h delay in the appearance of the roots (Fig. 3B). This suggests that *big* is inhibited in initiation, rather than emergence, of the WiR and is consistent with inhibition of auxin transport being responsible, at least in part, to the *big* root initiation phenotype.

### *ro/big* mutants are hypersensitive to Ca^2+^ in WiR formation

Application of exogenous auxin could not fully rescue *big* root initiation defects (Liu *et al*., 2022), prompting us to identify other factors that could be defective in *big*. The plant and mammalian *RO/BIG/UBR4* genes have a calmodulin-binding domain and it was suggested that *UBR4’s* function is to protect the integrity of subcellular organelles from increased Ca^2+^ levels during neuronal Ca^2+^ spikes (Belzil *et al*., 2013). In *Arabidopsis, big* mutants were reported to have increased cytosolic Ca^2+^accumulation in seedling leaves (Hearn *et al*., 2018), suggesting that it may be involved in Ca^2+^ homeostasis. As it was previously reported that Ca^2+^ levels can affect WiR initiation rates (Lanteri *et al*.,2006; Yu *et al*., 2019) we tested whether altered Ca^2+^ response may underlie some of the root initiation defects in *big*.

Treatment of tomato cuttings with 2 mM Ca^2+^ did not affect WT control plants but inhibited WiR root production in *ro* (Fig. 4A-B). Chelating the Ca^2+^ with EGTA restored root production (Fig. 4C-D), while the MgCl_2_ control treatment did not affect either WT or *ro* (Fig. 4E-F).

**Fig. 4.**
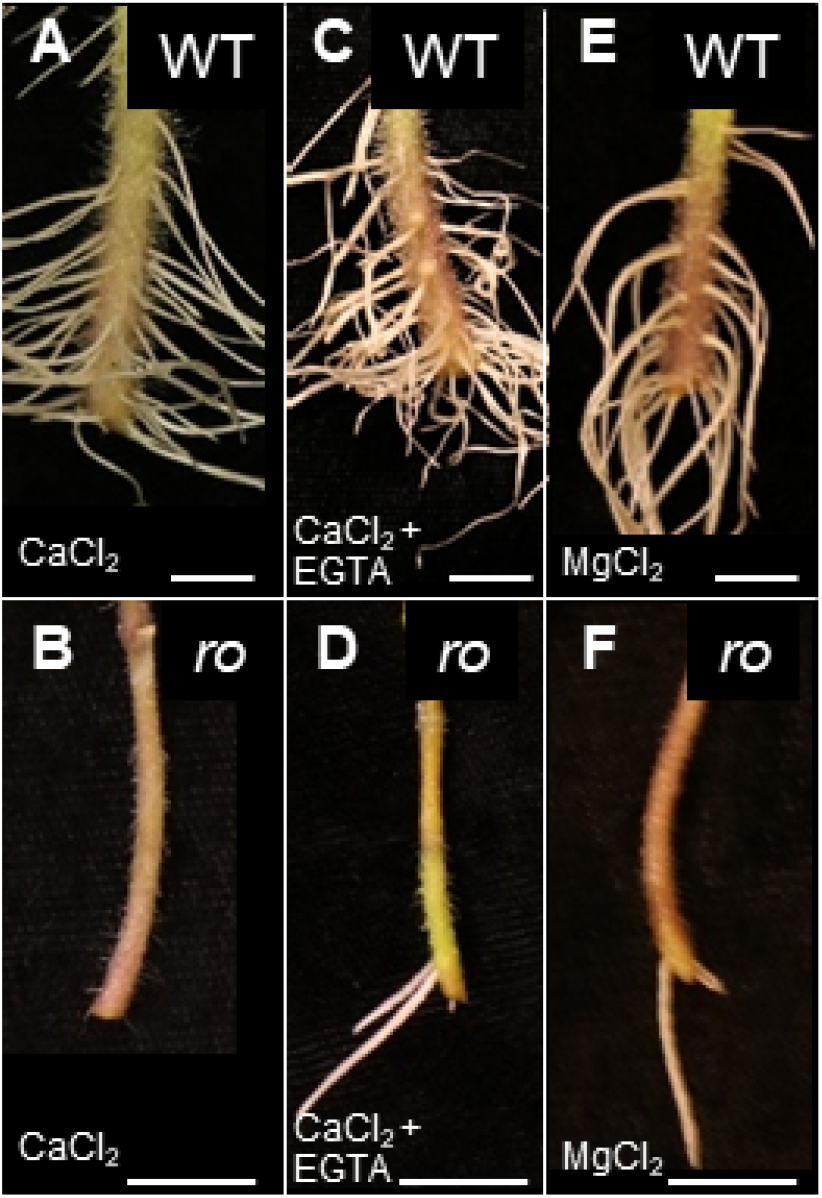
Root initiation in tomato *ro* mutant is sensitive to exogenous Ca^2+^. (A-F) One-month-old WT (A, C and E) or *ro* (B, D and F) tomato cuttings incubated in 2 mM CaCl_2_ (A and B), 2 mM CaCl_2_+ 2 mM EGTA (C and D) and 500 μM MgCl_2_ (E and F) for 7 days. Scale bars are 1 cm.

To test whether the sensitivity to calcium is a conserved trait of *ro/big* mutants, we tested the formation of WiR in *Arabidopsis* under mock conditions (no Ca^2+^) and with increasing concentrations of Ca^2+^. The number of hypocotyl-WiR was not affected by exogenous Ca^2+^ in WT *Arabidopsis* but was reduced in a dose-dependent manner in *big* mutants, suggesting that these mutants were sensitive to calcium also in *Arabidopsis* (Fig 5A). We note that higher concentrations of Ca^2+^ were required to observe an effect in *Arabidopsis*, as compared to tomato. Treatment with MgCl_2_ had the same effect on both WT and *big* mutants, suggesting that the hypersensitivity of the mutant is specific to Ca^2+^ (Fig. 5B). Consistently, a high level of Ca^2+^ had no effect on the *DR5rev:3xVenus-N7* signal in WT plants but delayed the appearance of the signal in *Arabidopsis big* mutants, where DR5 activation was not apparent even at 72h after the cut (Fig. 5C and compare to Fig. 3D). Overall, these results are consistent with altered Ca^2+^levels interfering with auxin transport in *ro/big* mutants.

**Fig. 5.**
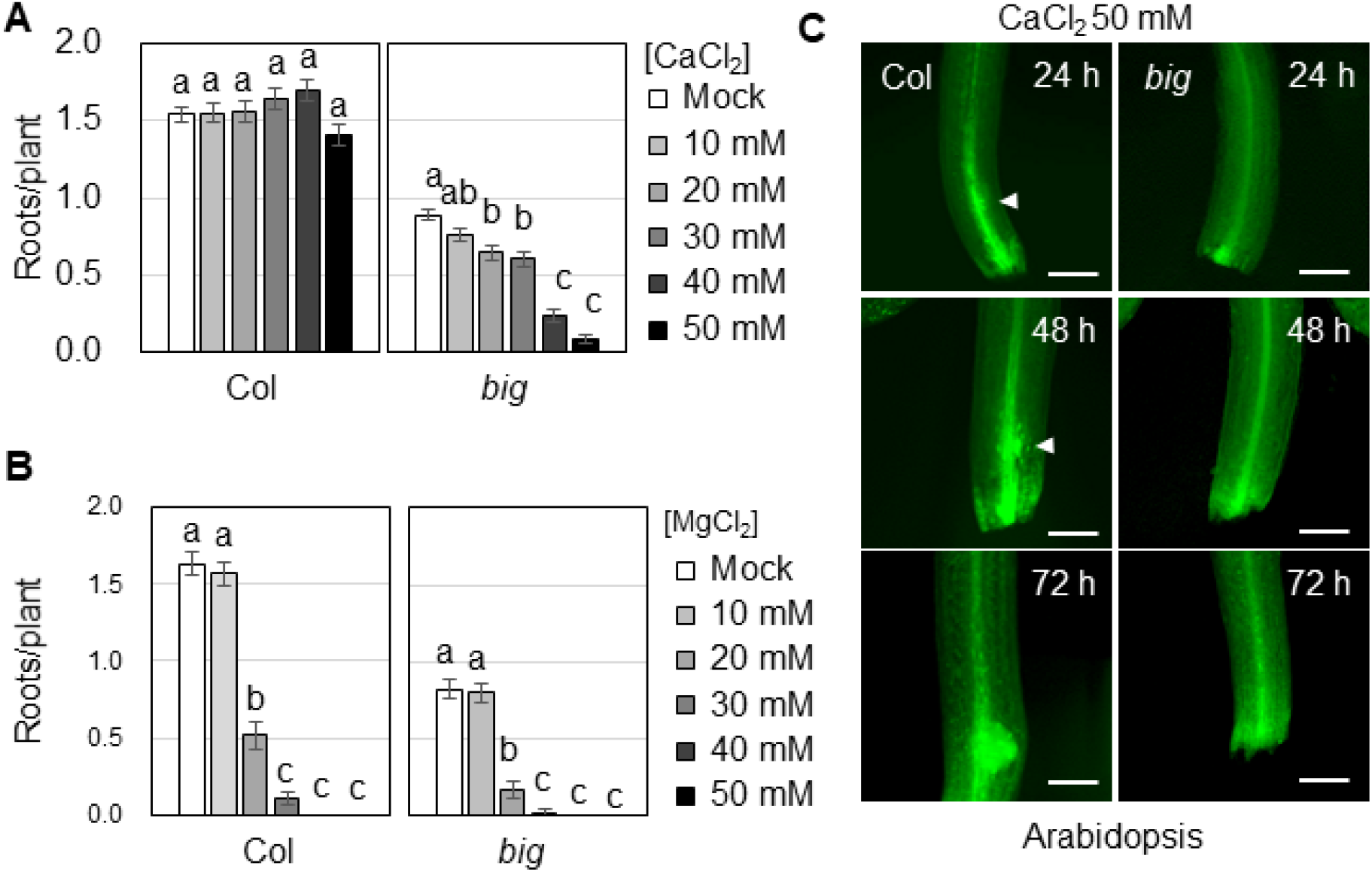
Arabidopsis *big* mutants are also sensitive to exogenous Ca^2+^. (A-B) Number of WiR of *Arabidopsis* Col and *big* mutant cuttings at 7 dac treated with different concentrations of CaCl_2_ (A) or MgCl_2_ (B) (n= 124, 106, 120, 114, 116 and 113 for Col and 89, 102, 116, 106, 106 and 105 for *big* in mock, 10, 20, 30, 40 and 50 mM CaCl_2_ respectively and n= 108, 112, 107, 121, 116 and 112 for Col and 91, 86, 96, 93, 103 and 104 *big* in mock, 10, 20, 30, 40 and 50 mM MgCl_2_ respectively). (C) Stereo microscope images of *DR5rev:3xVENUS-N7* cuttings incubated in 50 mM CaCl_2_ in WT (left) and *big* (right) backgrounds. Arrowheads mark root primordia. At least 6 plants were examined and representative images shown. Bars represent the mean ± SE; letters are statistically significant differences (p<0.01; ANOVA and Tukey’s post-hoc test). Scale bars are 200 μm.

### Ca^2+^ increases plant sensitivity to auxin transport inhibition

To determine whether the inhibitory effect of Ca^2+^ on WiR stems from a defect in the transport machinery, we co-treated WT and *big* plants with Ca^2+^ and IAA. The addition of 10 mM Ca^2+^, which by itself had little to no effect on root production in both WT and *big*, resulted in suppression of IAA promoting effect on root number (Fig. 6A). Treatment with the synthetic auxin 2,4-dichlorophenoxyacetic acid (2,4-D), which is not carried by the PIN proteins (Yang & Murphy, 2009) increased root production in both WT and *big* mutant cuttings but was not sensitive to calcium treatment. Moreover, 2,4-D had a significantly stronger effect on *big* mutants, increasing the number of roots 3.3-fold, compared to the 1.7-fold increase produced by IAA (Fig. 6B).

**Fig. 6.**
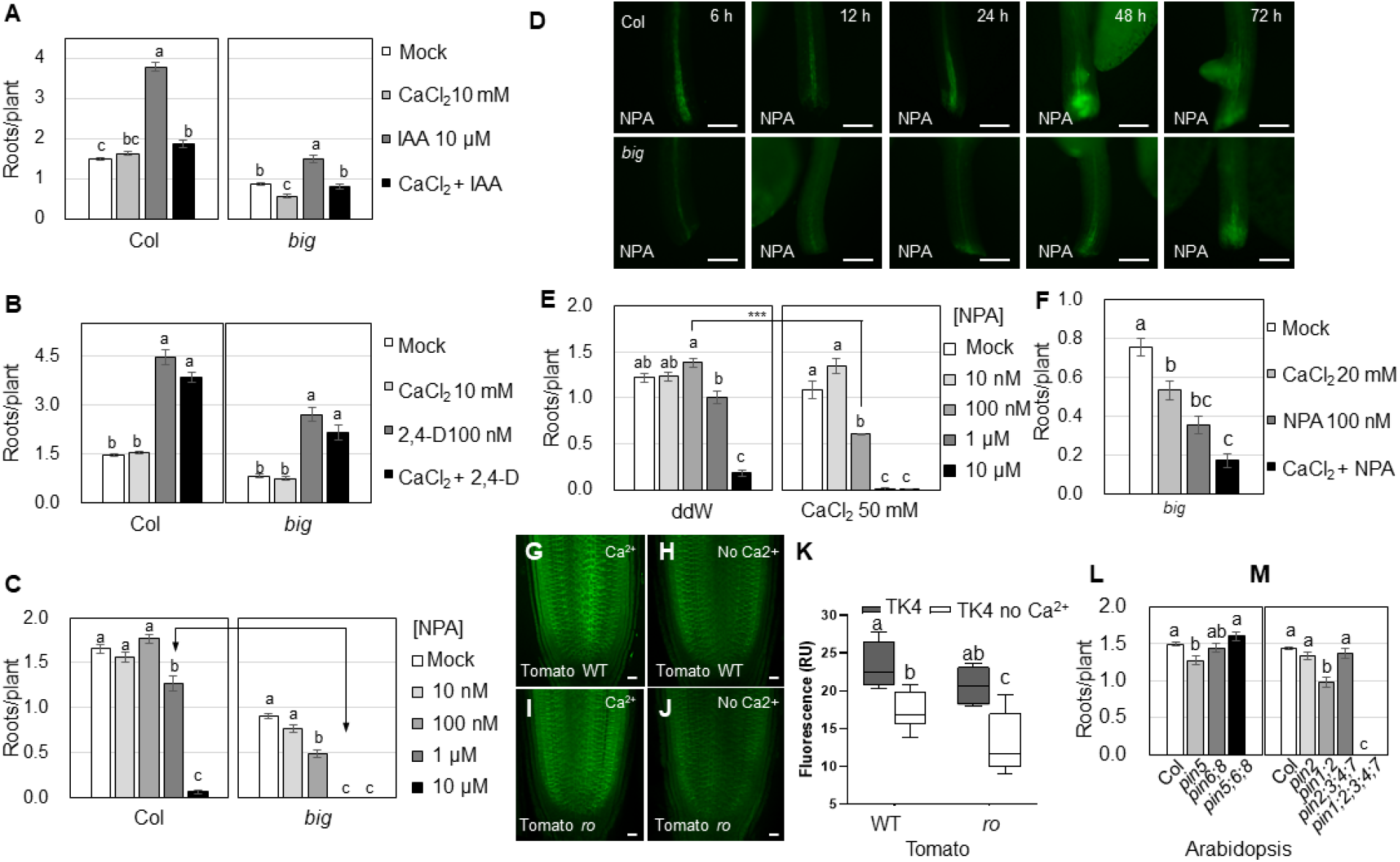
Altered Ca^2+^ levels control root initiation by inhibition of polar auxin transport. (A-C) WiR rates in 7 dac Col and *big* in different treatments (n= 125, 127, 133 and 130 for Col and 129, 129, 127 and 134 for *big* in mock, 10 mM CaCl_2_, 10 μM IAA and 10 mM CaCl_2_ + 10 μM IAA respectively; n= 120, 123, 108 and 115 for Col and 101, 94, 67 and 70 for *big* in mock, 10 mM CaCl_2_, 100 nM 2,4-D and 10 mM CaCl_2_ + 100 nM 2,4-D respectively; n= 119, 110, 124, 130 and 114 for Col and 119, 98, 131, 127 and 133 for *big* in mock, 10, 100 nM, 1 and 10 μM NPA respectively). (D) Stereo microscope images of *DR5rev:3xVENUS-N7* cuttings incubated in 100 nM NPA in WT (upper panels) and *big* (bottom panels) backgrounds. (E-F) Number of WiR in WT (E) and *big* (F) plants 7 dac cotreated with NPA and CaCl_2_ (WT, n= 116, 121, 129, 130 and 126 for no Ca^2+^ and 119, 119, 111, 115 and 111 for Ca^2+^ containing media in mock, 10, 100 nM, 1 and 10 μM NPA respectively; *big*, n= 119, 116, 113 and 121 for mock, 20 mM CaCl_2_, 100 nM NPA and 20 mM CaCl_2_ + 100 nM NPA respectively). (G-J) Confocal images of WT (G,H) and *ro* (I,J) tomato root meristems after 18h incubation with TK4 media (G,I) and calcium free TK4 media (H,J). (K) Quantification of fluorescence signals in (G-J), n=4. (L-M) Number of WiR in mutants of ER-PINS (L) and plasma membrane PINS (M) at 7 dac (n= 317, 98, 108 and 125 for Col, *pin5, pin6;8* and *pin5;6;8* respectively; n= 303, 84, 45, 106 and 84 for Col, *pin2, pin1;2, pin2;3;4;7* and *pin1;2;3;4;7* respectively). Bars represent the mean ± SE; letters are statistically significant differences (p<0.01; ANOVA and Tukey’s post-hoc test); Asterisks are statistically significant differences by two-tailed Student’s t-test (n.s. not significant, *** p<0.001). Scale bars are 200 μm in (D) and 20 μm (G-J).

If Ca^2+^ mediates its effect on root production by disruption of auxin transport, we can expect calcium treatment to act synergically with NPA and that *big* mutants would be sensitive to inhibition of auxin transport. In WT, treatment with 1 μM NPA resulted in a 23% reduction in root formation, while application of 10 μM NPA almost completely inhibited root initiation (96% reduction). *big* mutants were hypersensitive to NPA and a 46% reduction in root number was apparent already at 100 nM NPA. Unlike WT plants, 1 μM NPA was sufficient to completely block root initiation in *big* (Fig. 6C). NPA treatment prevented the activation of an auxin response signal near the cut in *big*, which is correlated with the lack of root initiation (Fig. 6D).

As expected from our hypothesis, treating cuttings with CaCl_2_ resulted in hypersensitivity to NPA. When CaCl_2_ was present in the media, WT plants exhibited a reduction in WiR number already at 100 nM NPA, and root production was abolished at 1 μM NPA, a response profile that phenocopied the response of *big* to NPA (Fig. 6C, E). Media containing calcium aggravated the response of *big* mutants to NPA (Fig. 6F). We note that *big* mutants were originally identified due to their resistance to NPA treatment in a primary root growth assay (Ruegger *et al*., 1997). The apparent contradiction between this and the NPA sensitivity shown here is probably due to the differences in development contexts (Gil *et al*., 2001).

Indeed, it was previously reported that NPA could induce the internalization of PIN proteins in *big* mutants, but not in WT (Gil *et al*., 2001). Further, levels of plasma membrane PINs in the root meristem were reduced in *big* mutants (Liu *et al*., 2022). We, therefore, tested whether the Ca^2+^ treatment could also affect PIN1 accumulation by using immunohistochemistry in the tomato root meristem. Supporting the link between Ca^2+^ and PIN localization, calcium depletion caused a reduction of *PIN1* immunostaining signal in the tomato root meristem. This reduction was much more pronounced in *ro* mutants (Fig. 6G-K). We note that in the root meristem, we observed a differential effect in low levels of exogenous Ca^2+^, while the phenotypic effect on root initiation was apparent at high Ca^2+^ levels. While we do not have a clear explanation for this discrepancy, the differential response suggests that these mutants are more sensitive to changes in Ca^2+^ levels than WT.

The auxin transport system is composed of plasma membrane and ER-bound PINs (Sauer & Kleine-Vehn, 2019) and both types were suggested to play a part in WiR (Xu *et al*., 2005; Simon *et al*., 2016; da Costa *et al*., 2020). To determine which PIN family plays the major role in WiR production, we tested WiR formation in single and high-order *pin* mutants. The effect of mutants in ER-localized PINs was low and changes in the number of roots were only significant in *pin5* mutant and with a mild 15% reduction in our experimental system. No aggravation of this phenotype was observed when all three ER-localized PINs were mutated, and their response was not significantly different from WT plants (Fig. 6L). In contrast, PM PINs were required for root initiation, with *pin1;2* mutants exhibited a 32% reduction in rooting rates and the quintuple mutant *pin1;2;3;4;7* not forming roots at all (Fig. 6M) indicating that the PM-PINs are the ones playing the major role in the regulation of WiR initiation in the *Arabidopsis* hypocotyl.

Taken together, these results suggest that in *ro/big* mutants, Ca^2+^ levels interact with polar auxin transport, possibly by disrupting plasma membrane PIN localization and preventing the auxin flow required for WiR initiation.

### BIG has ER localization signals and is required for maintaining cytoplasmic streaming

It is not clear how *RO/BIG* could control PIN activity since not much is known about the molecular function of this protein. In mammalians, UBR4/p600 was suggested to be localized to the ER where it acts to maintain ER integrity under high Ca^2+^ conditions (Shim *et al*., 2008; Belzil *et al*., 2013). To test whether such function may be conserved in plants, we first tested the intracellular localization of BIG. BIG is a 570 kDa protein and we were unable to construct a complete protein fusion. We, therefore, opted to generate partial protein fusions. We fused 6 parts of the protein to mScarlet-i and transiently expressed them in *Arabidopsis* cotyledons (Supplementary Table S1). Fusions for 5 of the 6 parts of the proteins did not produce any signal, suggesting that these are unstable. However, a fusion of the 251 aa C-terminal region resulted in ER-localization, which was verified by co-localizing with the ER marker YFP-HDEL (Fig. 7A-C).

**Fig. 7.**
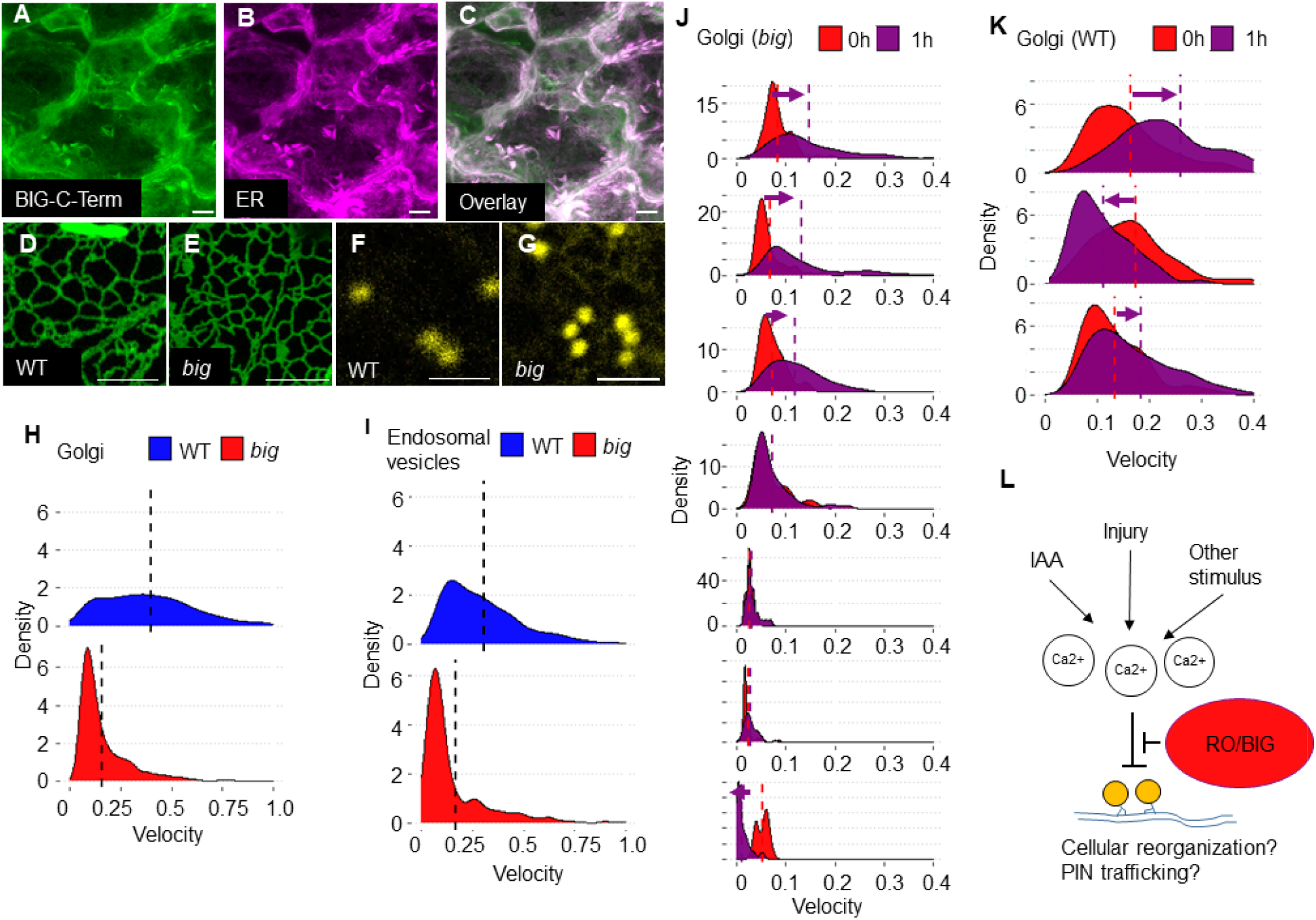
BIG is associated with the ER and is required for maintaining subcellular movement. (A-C) Maximal projection confocal images of the C-terminal region of the BIG protein fused to the mScarlet-i (A), the ER marker YFP-HYDEL (B) and the overlay of both (C) in epidermal cells of *Arabidopsis* cotyledons. (D-G) Confocal images of the ER marker GFP-HDEL (D and E) and the Golgi marker Man49-YFP (F and G) in Col (D and F) and *big* (E and G) epidermal cells of *Arabidopsis* cotyledons. (H-I) Distribution of Golgi bodies (H) and endosomal vesicles (J) velocity in Col and *big* mutant epidermal cells (Golgi, WT 0.4 ± 0.007 μm/s, big 0.16 ± 0.004 μm/s; n= 1123 and 954 for WT and *big* respectively; p<0.0001; Student’s t-test. endosomal vesicles, WT 0.31 ± 0.009 μm/s, big 0.17 ± 0.008 μm/s; n= 540 and 630 for WT and *big* respectively; p<0.0001; Student’s t-test). (J-K) Distribution of Golgi bodies velocity in individual epidermal cells of *big* (J) and WT control (K) at time 0 and 1 hour later. Each distribution plot represents a single cell observed at the two time points (Arrows mark a significant change in mean velocity between the timepoints, p<0.0001 in Student’s t-test), Dash lines indicate the mean velocity. (L) Hypothetical model for the function of RO/BIG. Scale bars are 10 μm.

To test whether BIG plays a role in maintaining the structure of intracellular organelles, we examined the morphology of the ER in *big* mutants using the ER-localized makers YFP-HDEL (Saint-Jore-Dupas *et al*.,2006; Nelson *et al*., 2007). Examination of cotyledon epidermal cells revealed no obvious structural disruption to the ER (Fig. 7D-E). However, we could observe abnormal inhibition of ER movement in cells of *big* mutants. To quantify this effect we used the Golgi marker Man49, as the Golgi is tethered to the ER and its punctate nature makes them easy to track (Sparkes *et al*., 2009). Golgi morphology, as evident by the Man49 marker appeared normal in *big* (Fig. 7F-G). Timelapse imaging of the Golgi in epidermal cells revealed a marked reduction in the average velocity of the Golgi organelle in the *Arabidopsis big* mutant. Notably, Golgi movement in 56% of the cells was almost completely arrested (Fig. 7H).

The cytoplasm of plant cells continuously undergoes intracellular flow, promoted by the activity of myosin motors. This flow is arrested when cytosolic Ca^2+^ concentrations are elevated (Tominaga & Ito, 2015) and enhanced in response to exogenous auxin (Friml *et al*., 2022). The observed arrest of Golgi and ER movement can be due to a specific effect on the ER or due to general arrest in cytoplasmic streaming and organelle movement. To distinguish between these possibilities, we measured the velocity of endosomal vesicles using the marker ARA6 (Ueda *et al*., 2001). The average velocity of endosomal vesicles was severely reduced in *big* mutants, to a similar extent as observed for the Golgi, with 53% of the cells exhibiting almost complete arrest of movement (Fig. 7I). To verify that this streaming arrest does not represent dead or dying cells, we performed a long-term timelapse recording of the Golgi marker in *big* epidermal cells (n=7) which appear arrested. One-hour-long recordings revealed that 3 out of the 7 cells resumed Golgi movement (Fig. 7J), indicating that the arrest of organelle movement is transient and that cells can resume their activity. Similar changes in rates of Golgi streaming were observed in WT cells (Fig. 7K), but the arrest of streaming was rare. Taken together, our data suggest that *big* mutants are defective in organelle movement within the cell, a function that is linked to the regulation of cytoplasmatic Ca^2+^ levels and short-term auxin response (Friml *et al*., 2022).

## Discussion

Many plant developmental processes rely on the accumulation of auxin at specific places in the plant. In response to cutting off the stem, changes to auxin transport lead to accumulation of auxin near the wound site which promotes the developmental response to wounding, such as root initiation. However, how injury and auxin transport are linked is unknown.

It is well documented that increased cytoplasmatic Ca^2+^ causes inhibition of intracellular movement, due to disruption of myosin activity (Tominaga & Ito, 2015) and it was shown that *big* mutants have high cytoplasmic Ca^2+^ concentrations (Hearn *et al*., 2018). Indeed, mutants in *big* have low rates of cytoplasmic streaming and can even exhibit temporary arrest of the movement, at least as measured in epidermal cells. Curiously, mutants in myosins XIs, which are defective in cytoplasmic stream, also exhibit defects in PIN1 polarization and reduced root initiation, similar to *big* (Peremyslov *et al*., 2015; Abu-Abied *et al*., 2018).

It was recently shown that auxin response enhances the rates of cytoplasmatic streaming and mutants which are defective in this response have lower rates of vasculature reconnection following injury, supporting the link between auxin response, cytoplasmic streaming, and downstream developmental responses (Friml *et al*., 2022). However, it is still unclear how and if Ca^2+^ levels and changes in cytoplasmatic streaming rates are translated to changes in development. One possibility is that defects in ER organization may lead to disruption to the activity of ER-localized PINs. Indeed, a previous study reported that mutants in the ER-localized *PIN6* have an increased rate of WiR (Simon *et al*., 2016). However, in our system, we could not observe any significant effect for the triple ER-PIN mutant *pin5;6;8* on WiR production. Rather, a mutant in all five PM-PINs could not form WiR, suggesting these are the key factors involved in channeling auxin to the site of root initiation. Thus, it is likely the function and organization of PM-PINs that needs to respond in other to produce roots in response to severe wounds.

In plants, both auxin and injury or touch were shown to increase cytoplasmic Ca^2+^ concentration (Vanneste & Friml, 2013; Kiep *et al*., 2015; Marhavý *et al*., 2019) and in the shoot meristem, these Ca^2+^ induced signals were important for reorientation of PM-PIN proteins (Li *et al*., 2019). Thus, we hypothesize a possible model where the role of RO/BIG is to buffer the increased Ca^2+^ concentration produced by auxin or other signals, allowing rapid cytoplasmic streaming and rapid deployment of PIN proteins to the membrane to alter auxin transport rates (Fig. 7L). This proposed role for RO/BIG is consistent with the suggested role for UBR4 in neurons, where it is important for buffering the deleterious effects caused by increased concentration of cytoplasmatic Ca^2+^ during neuronal activation (Belzil *et al*.,2013).

## Materials and Methods

### Plant materials and growth conditions

Tomato *(Solanum lycopersicum)* wild type (WT) are from M82 or Earliana cultivars. Seeds were obtained from the Tomato Genetics Resource Center (Davis). Tomato seeds were sown in soil and grown in long-day light conditions (16h:8h, 22°C) for four weeks and then transferred to a greenhouse and grown under natural light conditions. All *Arabidopsis thaliana* plants used in this study are in the Columbia (Col-0) background. *tir3-1* (Ruegger *et al*., 1997) plants were backcrossed to Col to segregate out the *gl1* mutation. Resequencing of the *tir3-1* allele indicated that the mutation is a C>T transition at position 9286 that causes a stop codon in the 8^th^ exon and not as previously reported (Supplementary Fig. S1E; (Gil *et al*., 2001). PIN mutants *pin1-1 eir1-1 (pin2), eir1-1 pin3-3 pin4-2 pin7En* (Verna *et al*.,2019), *pin1-1 eir1-1 (pin2) pin3-3 pin4-2 pin7En* (Verna *et al*., 2019), *pin6 pin8-1* (Sawchuk *et al*., 2013), *pin5-5* (Mravec *et al*., 2009) and *pin5-5pin6-2 pin8-1* (Le *et al*., 2014) and *DR5rev:3xVENUS-N7* (Heisler *et al*., 2005) were previously described.

### WiR assay

Roots from four-weeks-old tomato plants were completely removed by cutting 1-2 cm above the root-hypocotyl junction and resulting explants were incubated on rooting media; double distilled water (ddW), 10 μM IAA (Sigma I2886), 2 mM CaCl2, 2 mM CaCl2 + 2 mM EGTA (Sigma MFCD00004291), 500 μM MgCl_2_ in transparent flasks. The rooting solution was refreshed every two days. Experiments were performed in three or four replicates. For the rooting experiments, *Arabidopsis* seeds were sown on half-strength Murashige and Skoog medium, 2% agar plates, stratified for 3 days at 4°C in the dark, and then transferred to long-day light conditions (16h:8h, 22°C). Five-day-old *Arabidopsis* seedlings were cut above the root-hypocotyl junction to remove the root and the resulting explants were incubated on Petri dishes with Whatman paper and rooting solution (ddW, IAA, CaCl_2_, MgCl_2_, NPA (Chem Service N-12507), 2,4-D (Sigma D7299)). Between three and four independent replicates were done for each experiment, each one with at least 20 plants per genotype and condition.

### AR induction in cotyledon explants

Seeds were surface sterilized by immersion in 30% commercial bleach for 25 min followed by three rinses with sterile distilled water and aseptically sown in TK4 medium supplemented with 2% sucrose and 0.8% Agar in 90 mm Petri dishes. Cotyledons were excised from 7-8-day-old seedlings and placed with their abaxial surface in a TK4 medium containing 1.5 mM Ca^2+^ supplemented with 50 nM NAA in 90 mm Petri dishes. NAA was dissolved in DMSO and added to the medium after autoclaving to obtain a final DMSO concentration of less than 0.05%. Cotyledon explants were incubated in a growth chamber at 23 ± 1 °C with a photoperiod of 16 h light/8 h darkness and a photon fluence rate of 100 μmol m^-2^s^-1^.

### Cloning and transgenic plants

Binary plasmids were built through the Golden Gate cloning system as described in (Werner *et al*., 2012). *BIG gene* fragments I, II, IV, and V were synthesized by Syntezza Bioscience Ltd. in the pUC57-Kan backbone, and fragments III and VI were cloned in the pAGM1287 backbone (Primer for III are GTCAATCTCGGGAAAGACAAAG, CCCAGGCTTATTTCTTCGGGCTTTG and for VI: CGAGGTGGAGGAAGAGATAGC,). All BIG fragments were assembled as follow: 2×35s: *BIGfragment-* mScarlet-i:HSPterm in pICH47761. *Solyc02g089260* CRISPR binary plasmid was assembled as described in (Brooks *et al*., 2014) carrying two guides targeting the sequences GATACAAGATGCATGCTTCA and GTGGCTGTATCCACATTGTC in the N-region of the gene, the kanamycin resistance cassette from pICSL70004 and the CRISPR/cas9 cassette from pICH47742::2×35S-5’UTR-hCas9(STOP)-NOST. Tomato M82 transformation was performed according to (McCormick, 1991).

### IAA transport assay

Three hypocotyl sections (5 mm) from 7-week-old M82 and rosette (*ro*) mutant plants were used for basipetal IAA transport assays as described earlier (Nicolás *et al*., 2007; Cano *et al*., 2018). Briefly, these explants were placed in an upright position on a 1% agar block, and a 5 μl drop of a 200 μM labeled IAA ([^13^C]_6_C_4_H_9_NO_2_; OlChemIm, Olomuc, Czech Republic) solution was added to the upper cut end. Every 30 min and up to 240 min, the receiver agar block was collected and replaced by a new one. During the assay, explants were kept in darkness at 25° C. Ten μL of filtered extract were injected in a UPLC-HRMS system consisting of an Acquity UPLC (Waters Corporation, Mildford, MA, USA) coupled to a Xevo G2-XS QTOF mass spectrometer (Waters Corporation, Mildford, MA, USA) using an electrospray ionization (ESI) interface. Mass spectra were obtained using the MassLynx software version 4.2 (Waters Corporation, Mildford, MA, USA). For IAA quantification, a calibration curve was constructed (1, 10, 50, and 100 μg l^-1^). [^13^C_6_]-IAA was identified by extracting the exact mass from the full scan chromatogram obtained in the negative mode, adjusting a mass tolerance of ≤ 1 ppm. The concentrations of [^13^C_6_]-IAA were semi-quantitatively determined from the extracted area of the peak with the TargetLynx application manager (Waters Corporation, Mildford, MA, USA) by using the calibration curve of IAA. Each point on the curves corresponds to the mean value±SE. Linear traces of cumulative transported IAA were used to estimate transport intensity and velocity (Cano *et al*., 2018).

### Transient Expression on *Arabidopsis*

Three/four-day-old *Arabidopsis* seedlings were transformed with agrobacterium using the FAST technique (Li & Nebenführ, 2010) with the Golgi marker (Man49-YFP; (Nelson *et al*., 2007), the ER marker (GFP/YFP-HDEL; (Nelson *et al*., 2007), vesicles marker (ARA6-GFP; (Ueda *et al*., 2001) and/or *BIGfragment*-mScarlet-i.

### Imaging

Tomato hypocotyls were hand-sections using a razor, stained with SR2200 (0.1 % (v/v)), and mounted in water. *Arabidopsis* epidermal cotyledon cells were mounted in water. Tissue was observed using a Leica SP8 confocal microscope with ×20 dry or ×63 water objectives respectively. Lasers of 405, 488, and 552 nm were used for excitation of SR2200, GFP/YFP, and mScarlet-i, respectively. To visualize DR5 in Arabidopsis hypocotyls, tissue was cleared with ClearSee (Kurihara *et al*., 2015) and photographed using a Nikon SMZ18 fluorescence stereoscope with a 480nmEX/525nmEM filter.

### Immunohistochemistry

Roots from 6–9-day old seedlings were cut and incubated for 16 hours (overnight) in a TK4 liquid medium (control one with 1.5mM CaCl_2_ and without Ca^2+^). Roots were fixed and subjected to a standard immunolocalization procedure using PIN1 (7E7F) antibody (1:40 dilution) (incubation overnight with primary antibody at 4 °C and 1 h at 32 °C). Roots were mounted on slides with a double spacer (300 μm total thickness) and scanned with a LEICA Stellaris microscope (ex. 488, 515-550 emission) with a constant setting for all samples. The images were exported to tiff and the fluorescence was measured with the Fiji program.

### Measurement of vesicles and Golgi bodies movement

For analysis of cytoplasmic streaming, 20 to 60 seconds movies (≈1frame/s) of Arabidopsis cotyledons expressing the Man49-YFP Golgi marker or ARA6-GFP vesicles marker were recorded using an SP8 confocal microscope with 63x water lens. Timelapse images were examined to verify that tracked organelles are not moving out of plane. Only movies with at least 10 informative frames were used. For automatic extraction movement rates, movies were processed using TrackMate (Tinevez *et al*., 2017) in ImageJ software FIJI (Schindelin *et al*., 2012) using default parameters. Each movie resulted in 100-150 individual movement tracks. Density plots for movement rate distribution were generated using R 3.5.2.

## Data Availability

The data underlying this article are available in the article and in its online supplementary materials.

## Acknowledgments

We would like to thank Einat Sadot for critical reading of this manuscript, Jiri Friml for providing materials, and José Luis Micol for equipment.

