## Supplemental Figure 1 for "Mapping of the Classical Mutation *rosette* Highlights a Role for Calcium in Wound-induced Rooting"

Running head: Calcium in wound-induced rooting

Abelardo Modrego^1^, Moutasem Omary^1^, Alfonso Albacete^2^, Antonio Cano^3^, José Manuel Pérez-Pérez^4^ and Idan Efroni^1*^

^1^The Institute of Plant Sciences and Genetics in Agriculture, Faculty of Agriculture, The Hebrew University, Israel

^2^Institute for Agroenvironmental Research and Development of Murcia (IMIDA), c/Mayor s/n, 30150 La Alberca, Murcia, Spain

^3^Departamento de Biología Vegetal (Fisiología Vegetal), Universidad de Murcia, Murcia, Spain

^4^Instituto de Bioingeniería, Universidad Miguel Hernández, 03202 Elche, Spain

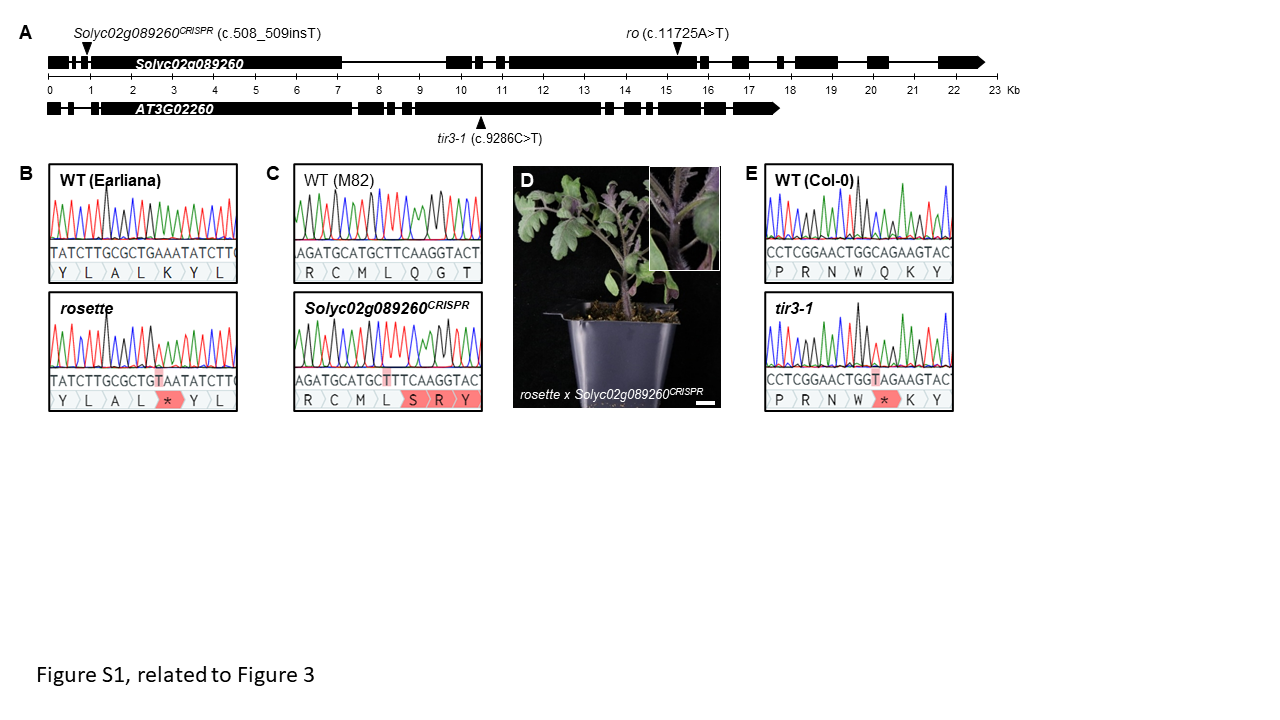

**Supplementary Figure S1. Mapping of the ro mutation**. (A) Gene structure of ROSETTE (Solyc02g089260) and BIG (AT3G02260), black boxes represent exons. Black triangles indicate the position of Solyc02g089260CRISPR, *ro*, and *big* mutations. (B-C) Chromatograms of the mutated region in Solyc02g089260 in Earliana and rosette (B), and M82 and Solyc02g089260CRISPR (C). (D) F1 progeny of a cross between ro and Solyc02g089260CRISPR has the ro phenotype. (E) Chromatograms of the region of AT3G02260 around position 9286 in WT (Col) and in tir3-1 allele of BIG.

Supplementary Table S1. ***BIG* gene fragments used to study the subcellular location of the protein**.

| Fragment | Coordinates (bp) | Domains |
| --- | --- | --- |
| I | from 1 to 3132 |  |
| II | from 3130 to 5225 | Zinc finger/UBR box |
| III | from 5222 to 8371 | WD40 domain |
| IV | from 8368 to 11518 | Zinc finger/U-box, Armadillo domain |
| V | from 11515 to 14541 | CaM binding domain, Armadillo domain,  E3 Ubiquitin Ligase UBR4 |
| VI | from 14539 to 15294 | E3 Ubiquitin Ligase UBR4 |
